# Designing a novel monitoring approach for the effects of space travel on astronauts’ health

**DOI:** 10.1101/2022.02.06.479323

**Authors:** Anurag Sakharkar, Jian Yang

## Abstract

Space exploration and extraterrestrial civilization have fascinated humankind since the earliest days of human history. However, it was only until last century that humankind finally began taking significant steps towards these goals by sending astronauts into space, landing on the moon, and building the International Space Station. However, space voyage is very challenging and dangerous, and astronauts are under constant space radiation and microgravity. It has been shown that astronauts are at a high risk of developing a broad range of diseases/disorders. Thus, it is critical to develop a rapid and effective assay to monitor astronauts’ health in space. In this study, gene expression and correlation patterns were analyzed for 10 astronauts (8 male and 2 female) using the publicly available microarray dataset E-GEOD-74708. We identified 218 differentially expressed genes between In-flight and Pre-flight and noticed that space travel decreased genome regulation and gene correlations across the entire genome, as well as individual signaling pathways. Furthermore, we rationally designed a rapid assay of 32 genes which could be used to monitor astronauts’ health during space travel. Further studies, including microgravity experiments, are warranted to optimize and validate the proposed assay.

## Introduction

“Space: the final frontier. These are the voyages of the starship *Enterprise*. Its continuing mission: to explore strange new worlds. To seek out new life and new civilizations. To boldly go where no one has gone before!”^1^. Every fan of the American science fiction TV series *Star Trek* is likely to remember this opening monologue. Indeed, space exploration and extraterrestrial civilization have fascinated humankind since the early days. However, it was only until last century that humankind finally took a solid step forward by sending astronauts into space and landing on the moon^2,3^. Into the 21^st^ century, humankind has made another great leap forward in space exploration, expediting the International Space Station (ISS) programme^4^, sending unmanned rovers to seek signs of life on Mars^5,6^, and starting private space ventures^7,8^. In addition, NASA (The National Aeronautics and Space Administration) is planning to send astronauts back to the moon in 2024 and to Mars in the 2030s^9^. Through all these endeavors, we have seen the twilight of manned space missions going beyond our harbor (Earth) and into deep space.

Space travel is challenging, dangerous and full of uncertainty. Astronauts are under sustained microgravity and cosmic radiation. Prolonged exposure to microgravity and cosmic radiation has been shown to cause a series of health problems, including loss of muscle mass^10,11^, reduced bone density^12,13^, compromised immune responses^14,15^, impaired renal functions^16^, neurological system irresponsiveness^17,18^, and development of cardiovascular diseases^19,20^. Furthermore, microgravity and cosmic radiation can cause several types of cancer, such as leukemia, likely due to compromised immunity^21^. However, studies under simulated microgravity also show that microgravity affects cell proliferation and apoptosis and can change cancer cells into less malignant phenotypes^22-24^. These studies indicate that the impact of microgravity and cosmic radiation on human health is profound, and it is unlikely that this impact is due to changes in one gene or a small group of genes. Therefore, genomic, transcriptomic and/or proteomic studies are crucial to fully understand the effects of microgravity and cosmic radiation on human health. These studies can lead to the development of effective strategies to monitor, prevent, and/or alleviate the effects on human health caused by space travel.

Approximately 12 years ago, NASA and JAXA (Japan Aerospace Exploration Agency) undertook a microarray study on 10 astronauts (8 male and 2 female), who had a six-month mission at the ISS^25^. Terada *et al*. used qPCR to confirm that space flight changed gene expression in astronauts^25^. Several genes, including *FGF18, ANGPTL7* and *COMP*, were upregulated and might play a role in inhibiting cell proliferation in hair follicles. However, they neither discussed the potential effects of the differentially expressed genes on the astronauts’ health nor provided any strategy to monitor these effects. Thus, in the current study, we reanalyzed this microarray dataset, rationally developed a method which could monitor the effects of space travel on astronauts’ health and searched for potential intervention options.

## Results and discussion

### Differentially expressed genes (DEGs)

NASA and JAXA undertook Study GLDS-174: “Effects of a closed space environment on gene expression in hair follicles of astronauts in the International Space Station” between 2009 and 2013. However, Terada *et al*. only reported the expression of selected genes in this study and did not undertake further analysis^25^. To better understand how space travel affects astronauts’ health, we downloaded the microarray dataset (E-GEOD-74708) from the NASA GeneLab open data repository. The microarray dataset includes gene expression information for three stages: Pre-flight (6 months to 2 weeks before launch), In-flight (while staying in the ISS), and Post-flight (2 days to 3 months after returning from the ISS). We then analyzed the dataset for gene expression and gene pair correlation. Using cut-off criteria of |log_2_FC| ≥ 3.00 and *p* < 0.05, 218 DEGs were identified between In-flight and Pre-flight (Supplementary File 1: DEG_list.xls). The top 20 up-and down-regulated genes are summarized in Table 1. However, no DEGs were detected between Post-flight and In-flight, suggesting that the astronauts may need a longer time than current guidelines for the body to adjust to the ground environment. Additionally, astronauts have a much higher risk of developing various types of diseases than normal people and precautionary measures are critical to protect their health after retuning to ground.

**Table 1.** Top 20 up- and down-regulated DEGs (differentially expressed genes) between In-flight and Pre-flight. The cut-off criteria for DEGs are |log_2_FC| ≥ 3.00 and *p* < 0.05.

| UP-REGULATED GENES |  |  | DOWN-REGULATED GENES |  |  |
| --- | --- | --- | --- | --- | --- |
| Gene Name | Log<sub>2</sub>FC | *p* Value | Gene Name | Log<sub>2</sub>FC | *p* Value |
| *LIMCH1* | 7.59 | 0.03 | *NFATC1* | -10.40 | 0.00 |
| *IER3* | 7.45 | 0.03 | *KIDINS220* | -10.35 | 0.03 |
| *ZNF664* | 6.57 | 0.03 | *ZCCHC9* | -7.82 | 0.00 |
| *NDUFA1* | 6.27 | 0.03 | *IGSF9* | -6.08 | 0.03 |
| *AL391650.1* | 5.28 | 0.03 | *HSD11B1L* | -5.76 | 0.00 |
| *TUBGCP5* | 5.07 | 0.03 | *CLTB* | -5.69 | 0.00 |
| *GAPVD1* | 5.03 | 0.03 | *YIPF2* | -5.68 | 0.03 |
| *AARS2* | 4.81 | 0.03 | *BTBD9* | -5.49 | 0.00 |
| *AC004080.5* | 4.77 | 0.03 | *LINC00668* | -5.41 | 0.00 |
| *NAA60* | 4.72 | 0.03 | *AL096711.2* | -4.97 | 0.03 |
| *IGBP1P1* | 4.41 | 0.03 | *EIF4E2* | -4.97 | 0.03 |
| *FANCD2* | 4.28 | 0.03 | *SHE* | -4.41 | 0.03 |
| *AMT* | 4.20 | 0.03 | *HOXC4* | -4.27 | 0.00 |
| *PNPLA4* | 4.11 | 0.03 | *THBS3* | -4.17 | 0.00 |
| *RABGAP1* | 4.10 | 0.03 | *PGM2L1* | -3.95 | 0.00 |
| *MAPKAPK5* | 3.89 | 0.03 | *TCEANC* | -3.93 | 0.03 |
| *DFFBP1* | 3.83 | 0.03 | *ARHGAP9* | -3.91 | 0.00 |
| *STAG1* | 3.71 | 0.03 | *ZNF451* | -3.80 | 0.03 |
| *SPON2* | 3.62 | 0.04 | *NCEH1* | -3.73 | 0.00 |
| *CHN1* | 3.59 | 0.04 | *AC010531.1* | -3.71 | 0.03 |

As expected, the DEGs between In-flight and Pre-flight regulate a broad spectrum of biological and physiological functions. For example, the most up-regulated gene *LIMCH1*, encoding LIM and calponin-homology domains 1, activates the non-muscle myosin IIa complex, stabilizes focal adhesion, and inhibits cell migration^26,27^. Thus, the upregulation of *LIMCH1* may explain observations that microgravity exposure can change cancer cells into less malignant phenotypes^22-24^. The second most up-regulated gene *IER3* encodes Immediate Early Response 3 and regulates cell apoptosis^28^, imflammation^29^ and tumorigenesis^30^. Upregulation of *IER3* has been observed in different types of cancer and may regulate cancer progression^31,32^. This implies another possible reason, other than radiation exposure, which increases the risk of developing cancer in astronauts during a long period of space travel. The most down-regulated gene, *NFATC1*, encodes Nuclear Factor of Activated T Cells 1 and plays important roles in osteoblast differentiation^33^, osteoclastogenesis^34^, T-cell differentiation^35^, lymphatic endothelial development^36^, cardiac valve morphogenesis^37^ and tumorigenesis^38^. The downregulation of *NFATC1* may result in osteoporosis, immunocompromised conditions, and prostate cancer in astronauts during a prolonged space stay. The second most down-regulated gene, *KIDINS220*, encoding Kinase D Interacting Substrate 220, modulates the development and function of the nerve and cardiovascular systems^39,40^. The downregulation of *KIDINS220* may contribute to the development of neurological disorders and cardiovascular diseases in astronauts. In summary, the large variation of the biological processes regulated by the DEGs provides a valuable resource of genes for us to develop a rapid assay kit (∼20-30 genes) to monitor astronauts’ health conditions in space. However, it is noteworthy that the three genes reported by Terada *et al*., *FGF18, ANGPTL7* and *COMP*, are not in the aforementioned DEG list as they failed to meet the cut-off criteria.

### Gene pair correlations

As mentioned above, the effect of space travel on astronaut’s health is comprehensive and involves the regulation of many genes. It is also well-known that any biological, physiological, or pathophysiological change requires a delicate coordination of multiple genes. Thus, we decided to carry out a gene pair correlation analysis for the whole genome (30645 transcripts for 23115 genes) among the Pre-flight, In-flight, and Post-flight expression profiles. As shown in Figure 1A, the genes are highly correlated in the Pre-flight, which This is consistent with previous studies showing that gene expressions are correlated in normal human tissues^41^. Upon staying in the ISS, the genome appeared to significantly lose its regulation of the gene expression and gene coordination was scrambled (Figure 1B). Our previous studies have shown that loss of gene pair correlations is a hallmark of carcinogenesis and/or cancer progression^42-44^. Therefore, loss of coordination of gene expression, other than cosmic radiation and compromised immune system, is likely to be another factor which increases the risk of cancer development in astronauts. Analysis of the Post-flight dataset showed that returning to ground did not significantly improve genome regulation and coordination of gene expression (Figure 1C), implicating that the effect of space travel on human health is more profound and longer lasting than currently thought. Extended care is necessary for astronauts to minimize the risk of disease development. As we progress towards manned missions into outer space (for example, to the Moon and Mars) and other prolonged space stays, it is critical to develop rapid approaches to monitor genome regulation and establish corresponding protocols to alleviate the effects of space travel on human health. Another important question that needs to be addressed is whether the human body could establish a new “routine” for prolonged space stays and what this would look like in terms of astronauts’ health. We speculate that loss of gene expression coordination might also be responsible for other diseases/conditions astronauts experience, such as muscle loss and compromised immune system. Extensive further studies in astrobiology, astromedicine and astropharmacy are warranted to prepare us for longer-term outer space exploration.

**Figure 1.**
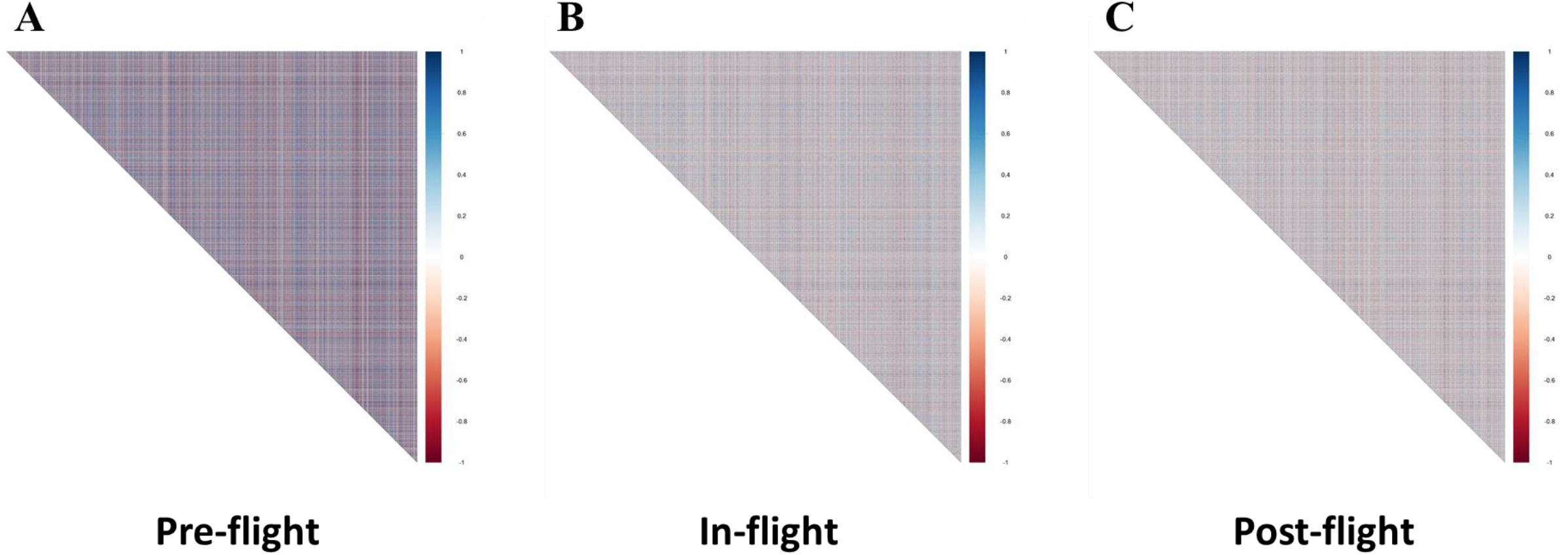
Gene pair correlations of the whole genome (30645 transcripts for 23115 genes) for 10 astronauts during Pre-flight (**A**), In-flight (**B**) and Post-flight (**C**). Positive and negative correlations are represented by blue and red, respectively.

### Signaling pathway and disease network of the DEGs

Since DEGs are the most altered genes during a biological process, we conducted a signaling pathway analysis of the 218 DEGs between In-flight and Pre-flight to figure out what biological functions are regulated by these genes. The top 11 signaling pathways were identified to be signal transduction (16 genes: *ARHGAP9, CCL2, CCNC, CHN1, CLTB, COL4A4, CREB1, CRHR1, CTNNBIP1, HIF1A, KIDINS220, NFATC1, PDPK1, SOS2, THBS3* and *YES1*), immune system (11 genes: *ATF2, BIRC2, CREB1, EIF4E2, IL7, NFATC1, PDPK1, UBA5, UBR4, XAF1* and *YES1*), gene expression (10 genes: *AARS2, CCNC, RRN3, ZNF184, ZNF253, ZNF529, ZNF606, ZNF664, ZNF699* and *ZNF711*), metabolism (10 genes: *ACSL4, ARSK, CCNC, GM2A, GPT, HACL1, NDUFA1, PIKFYVE* and *PSAT1*), metabolism of proteins (9 genes: *ARSK, CCL2, DPP4, GNE, MAGT1, PCSK1, SPON2* and *XRN2*), generic transcription pathway (8 genes: *CCNC, ZNF184, ZNF253, ZNF529, ZNF606, ZNF664, ZNF699* and *ZNF711*), developmental biology (7 genes: *CCNC, CLTB, COL4A4, CREB1, SCN2B, SOS2* and *YES1*), metabolism of lipids and lipoproteins (6 genes: *ACSL4, ARSK, CCNC, GM2A, HACL1* and *PIKFYVE*), axon guidance (6 genes: *CLTB, COL4A4, CREB1, SCN2B, SOS2* and *YES1*), innate immune system (6 genes: *ATF2, BIRC2, CREB1, NFATC1, PDPK1* and *YES1*) and disease (6 genes: *CCNC, CHMP4C, CREB1, CTNNBIP1, HIF1A* and *PDPK1*). In total, 48 genes were found to be involved in the regulation of these top 11 signaling pathways. Moreover, it is obvious that most of these DEGs are involved in the regulation of multiple signaling pathways, and that these signaling pathways regulate a broad range of biological, physiological and/or pathophysiological processes. This analysis reconfirms that space travel imposes comprehensive effects on the human system rather than affecting an individual organ or tissue.

We further analyzed the disease network for the 218 DEGs between the In-flight and Pre-flight expression profiles to identify the major disease/disorder conditions associated with space travel (Supplementary File 2: Disease_DEG.xls). As shown in Table 2, the top 20 disease/disorder conditions can broadly be divided into three categories: neoplasia/carcinoma, neurological disorder, and liver function, with the top 3 conditions being malignant neoplasm of the breast, colorectal carcinoma and malignant neoplasm of the prostate. More DEGs were associated with tumor development than other diseases/disorders. In 2019, Reynolds *et al*. reported the effect of space radiation on astronauts’ death using a statistical analysis^45^. Of the astronauts and cosmonauts who traveled to space between 1960 and 2018, 53 NASA astronauts have died, with 16 of those deaths (30.2%) being caused by cancer. Nevertheless, their study indicates that space radiation does not have a strong impact on the mortality of astronauts. This implies that the weakened genome regulation (*i*.*e*., reduced gene expression correlation) we observed in this study, other than compromised immune function, is likely to play an important role in tumor development in astronauts, even though the tumors may be less malignant or even benign. Moreover, consistent with previous studies^46,47^, our results showed that space travel significantly affects astronauts’ liver function and liver metabolism. Key genes identified from our analysis, such as *GPT* and *UBA5*, might serve as valuable monitoring biomarkers of astronauts’ liver function and even possible medical intervention points to protect astronauts’ health. Since liver is a major organ for drug metabolism, a systematic astropharmacy study is needed to establish drug profiles, including dosing, ADMET (absorption, distribution, metabolism, excretion, and toxicity) and even formulation, in preparation for future voyages into outer space. Space drug usage based on current information may be less effective or even detrimental to astronauts’ health.

**Table 2.** Top 20 disease/disorder conditions and their respectively associated DEGs based on disease network analysis of 218 DEGs between In-flight and Pre-flight.

| **DISEASE/DISORDER** | **GENES** |
| --- | --- |
| Malignant Neoplasm of Breast | *THBS3, UBR4, ATF2, NBN, CRHR1, AREG, HIF1A, PDPK1, COL7A1, ZNF404, BIRC2* |
| Colorectal Carcinoma | *POSTN, SACS, FANCG, XAF1, ACSL4, INTS13, COL7A1, NFATC1, C12ORF76, NDUFA1* |
| Malignant Neoplasm of Prostate | *GREB1, HMGN5, SPON2, HIF1A, CRYL1, CASZ1, ACSL4, NBN* |
| Prostatic Neoplasms | *CASZ1, NBN, GREB1, HIF1A, CRYL1, ACSL4, SPON2, HMGN5* |
| Schizophrenia | *PSAT1, BTBD9, CFAP65, CREB1, CCL2, HSPA12A, DKK3, VRK2* |
| Breast Carcinoma | *BIRC2, COL7A1, ZNF404, HIF1A, AREG, CRHR1, PDPK1* |
| Mammary Carcinoma, Human | *AREG, COL7A1, HIF1A, CRHR1, ZNF404, PDPK1, BIRC2* |
| Mammary Neoplasms | *AREG, COL7A1, PDPK1, ZNF404, CRHR1, HIF1A, BIRC2* |
| Mammary Neoplasms, Human | *ZNF404, COL7A1, PDPK1, CRHR1, BIRC2, HIF1A, AREG* |
| Unipolar Depression | *CCL2, PEA15, HIF1A, CRHR1, CREB1, ACSL4* |
| Liver Cirrhosis, Experimental | *GPT, TM6SF1, SGCB, ARHGAP9, CCL2* |
| Major Depressive Disorder | *HIF1A, CRHR1, PEA15, CCL2, CREB1* |
| Non-small Cell Lung Carcinoma | *E2F8, PSAT1, AREG, HIF1A, COL7A1* |
| Bipolar Disorder | *HIF1A, CRHR1, CREB1, HMGXB4* |
| Chemical And Drug Induced Liver Injury | *HACL1, GPT, UBA5, CCL2* |
| Chemical-induced Liver Toxicity | *HACL1, UBA5, GPT, CCL2* |
| Depressive Disorder | *CREB1, CRHR1, DPP4, ACSL4* |
| Disease Exacerbation | *COL7A1, E2F8, ATF2, HIF1A* |
| Drug-induced Acute Liver Injury | *GPT, HACL1, CCL2, UBA5* |
| Drug-induced Liver Disease | *HACL1, GPT, CCL2, UBA5* |

### Design of a rapid assay for space travel

Any biological or physiological process requires a delicate regulation of genes, including gene expression levels, interactions, and correlations. However, most genomics/genetic studies are focused on gene expression level rather than gene pair correlation. Although gene co-expressions and protein co-localizations are commonly studied in biomedical research, our research laboratory, to the best of our knowledge, is the first to apply gene pair correlation coefficient (a mathematical term) to describe and explain biological and medical questions^42-44^. Thus, in order to design a rapid assay to monitor astronauts’ health, we decided to adopt a set of genes with not only expression levels but also gene pair correlations significantly altered by space travel.

We calculated gene pair correlation coefficients (designated as CC) of the DEGs associated with the top 11 signaling pathways for the Pre-flight, In-flight and Post-flight datasets, and analyzed the gene pair correlation coefficient differentials (designated as ΔCC) between the In-flight and Pre-flight and between the Post-flight and In-flight (Figure 2). The changes in gene pair CCs between the In-flight and Pre-flight datasets were obvious for every single signaling pathway except the generic transcription pathway. However, the changes between Post-flight and In-flight CCs were minimal. Thus, using |ΔCC| > 0.70 as a cut-off, we identified gene pairs with their correlation coefficients significantly altered between the In-flight and Pre-flight datasets and summarized them in Table 3. These gene pairs were then classified into four categories based on changes in correlation coefficients: Category 1 – positive to more/less positive (highlighted in blue), Category 2 – positive to negative (highlighted in green), Category 3 – negative to positive (highlighted in brown), and Category 4 – negative to more/less negative (highlighted in red). It is noteworthy that Categories 2 and 3 contained more gene pairs than Categories 1 and 4, implying that many genes lost their coordination and even started counteractive processes in space. This observation supports previous studies that spaceflight significantly affects gene expression and homeostasis^48^. Apparently, this dramatic change in gene expression coordination affects normal biological and physiological functions in the human body and is likely detrimental to astronauts’ health.

**Figure 2.**
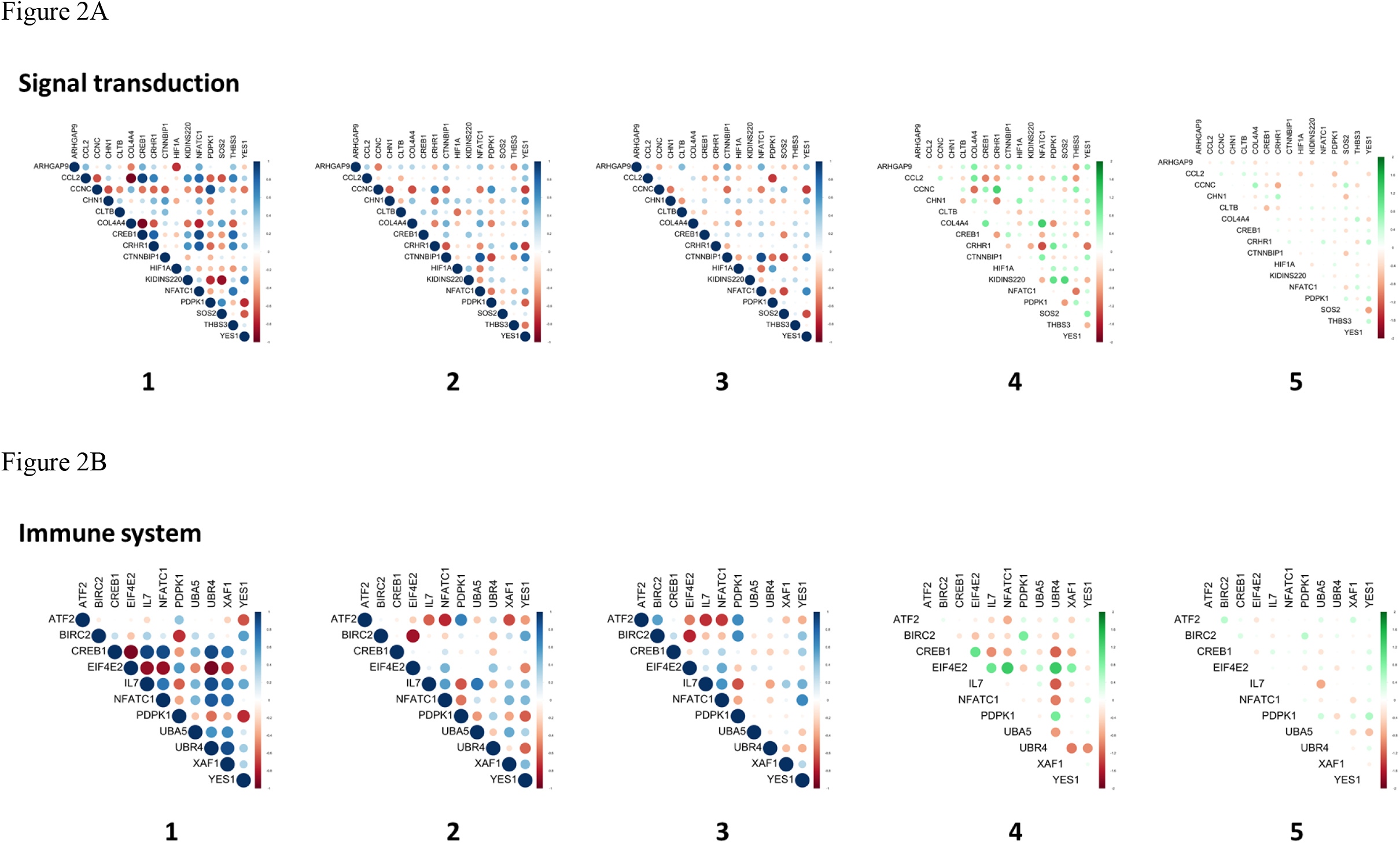
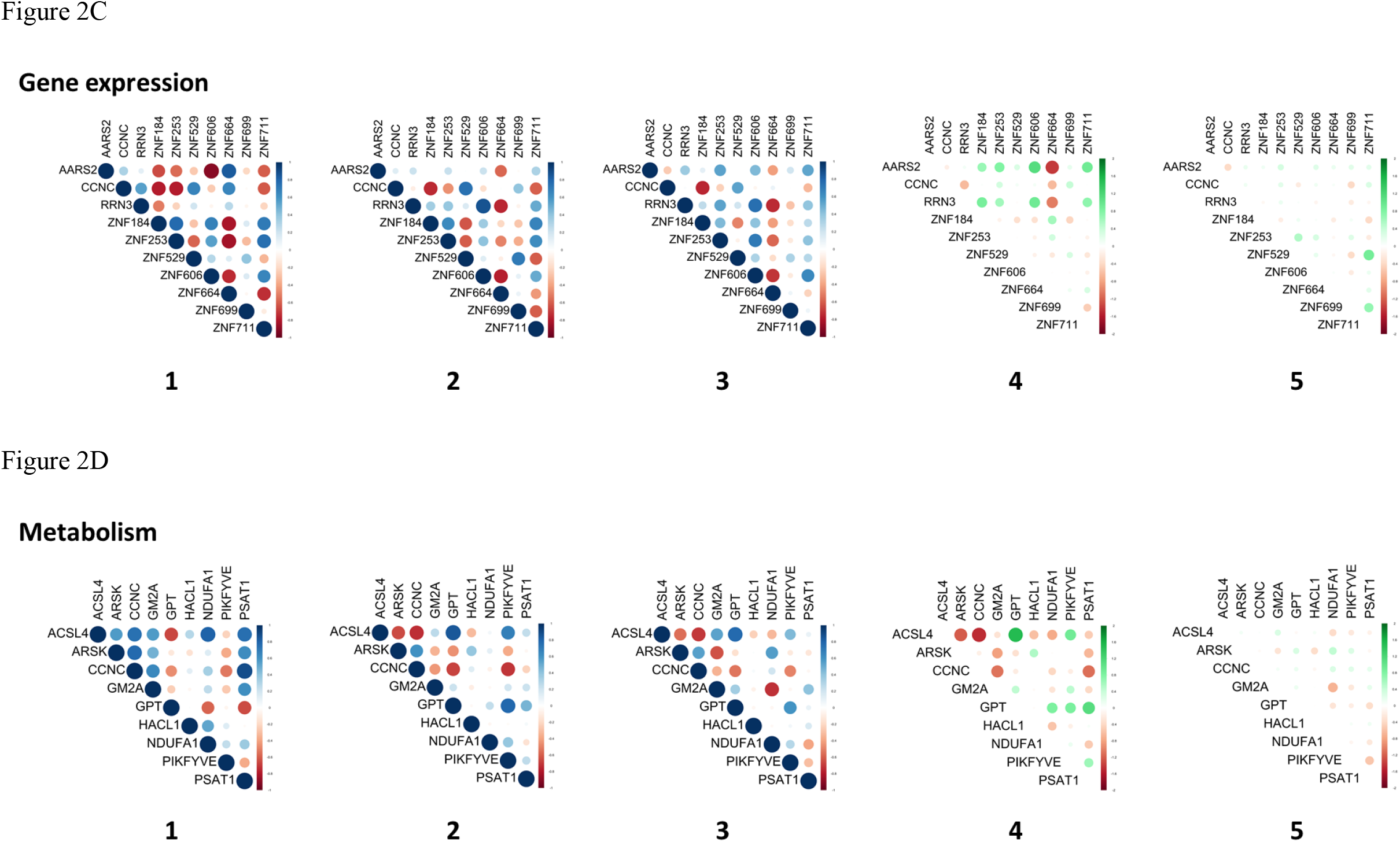
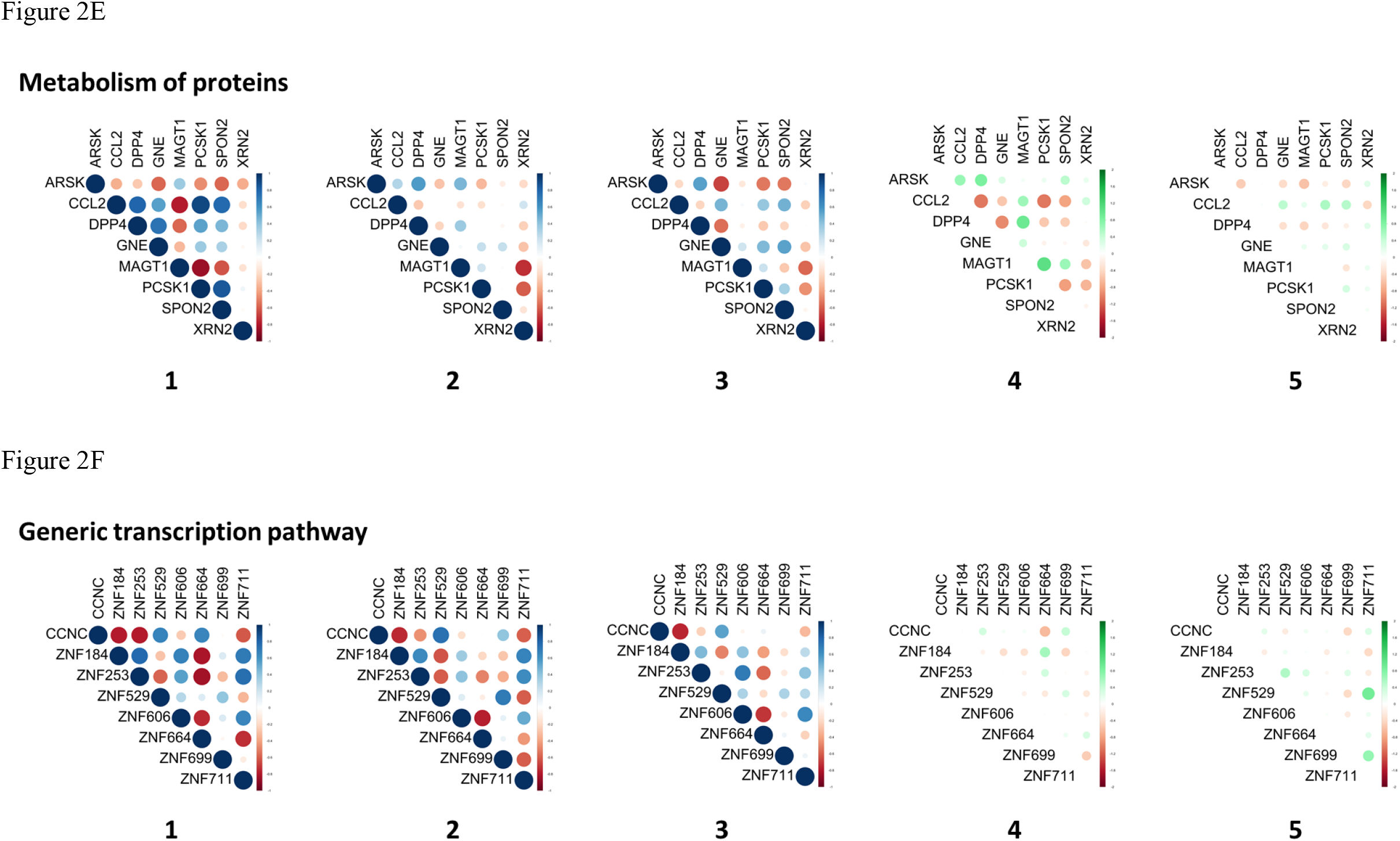
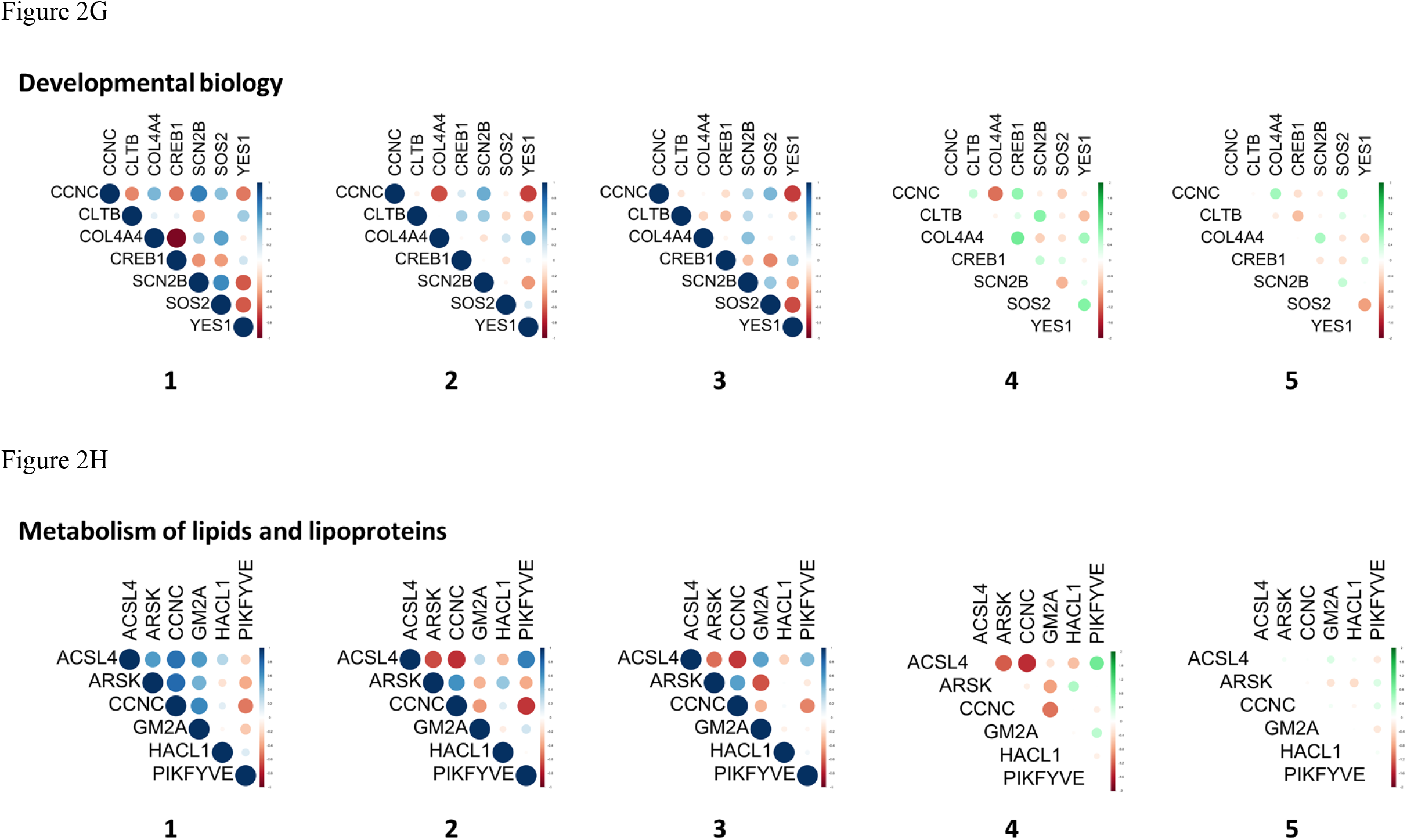
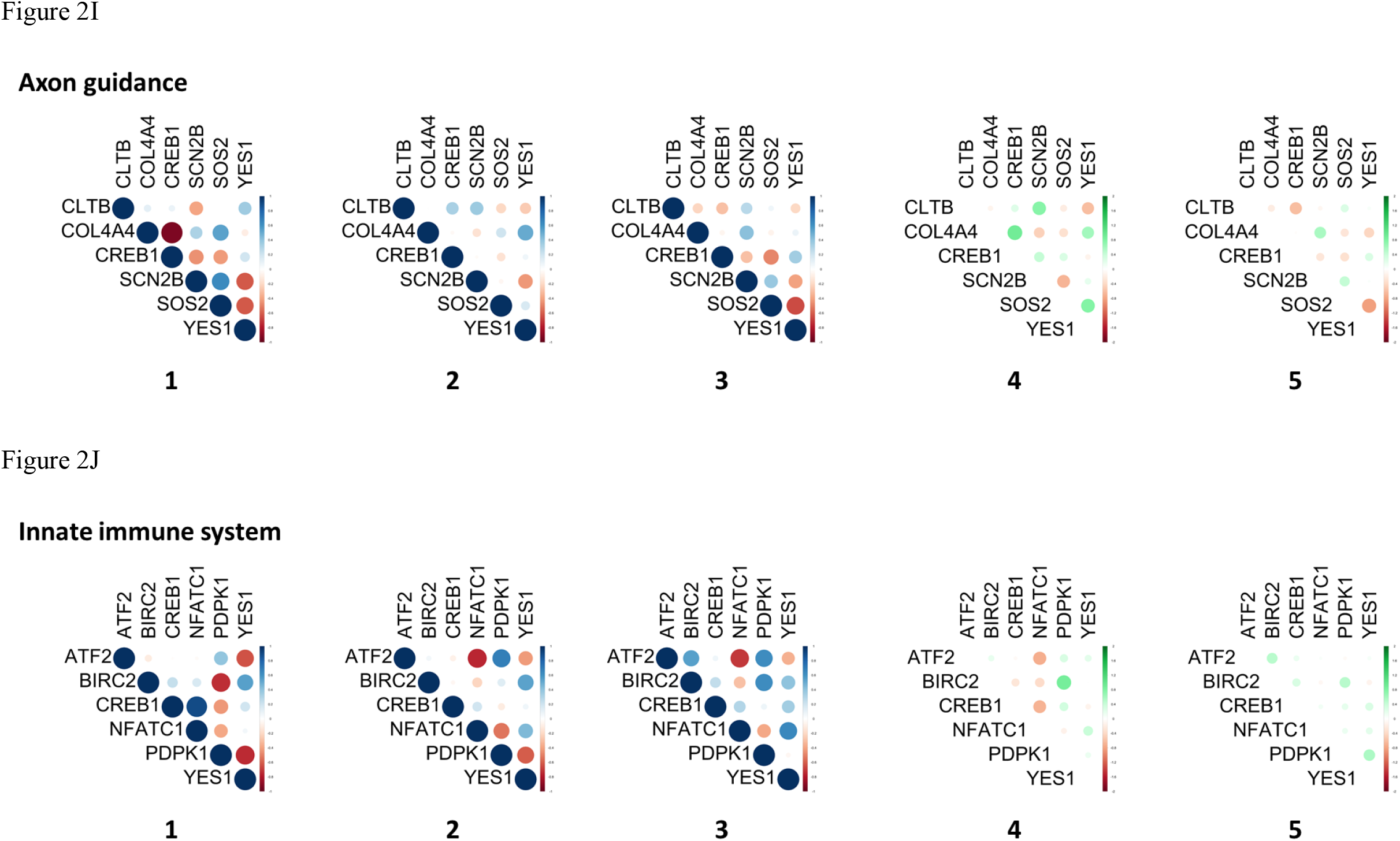
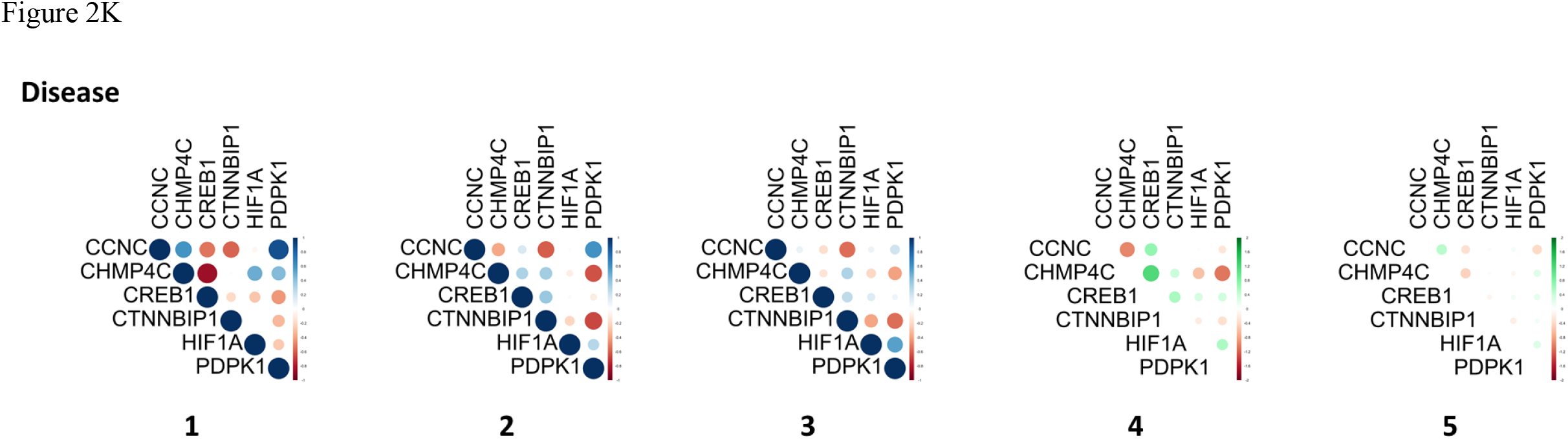
Gene pair correlations (**1**: Pre-flight, **2**: In-flight, and **3**: Post-flight) and differentials of gene pair correlations (**4**: between In-flight and Pre-flight, and **5**: between Post-flight and In-flight) for the differentially expressed genes (DEGs) in the top 11 signaling pathways: signal transduction (**A**), immune system (**B**), gene expression (**C**), metabolism (**D**), metabolism of proteins (**E**), generic transcription pathway (**F**), developmental biology (**G**), metabolism of lipids and lipoproteins (**H**), axon guidance (I), innate immune system (**J**) and disease (**K**). The top 11 signaling pathways were identified from InnateDB analysis of the 218 DEGs (|log_2_FC| ≥ 3.00 and *p* < 0.05). Positive and negative gene pair correlations are represented in blue and red, respectively; and positive and negative differentials of gene pair correlations are represented in green and red, respectively.

**Table 3.** Gene pairs with significantly altered correlation coefficients between In-flight and Pre-flight (i.e., |ΔCC (CC_In-flight_ – CC_Pre-flight_)| > 0.70) in the top 11 signaling pathways identified from signaling pathway analysis of the 218 DEGs between In-flight and Pre-flight. Gene pairs with correlation coefficients changed from positive to less/more positive (Category 1), positive to negative (Category 2), negative to positive (Category 3) and negative to less/more negative (Category 4) were highlighted in blue, green, brown, and red, respectively.

| GENE PAIRS | CC<sub>Pre-flight</sub> | CC<sub>In-flight</sub> | $\Delta\text{CC}$ |
| --- | --- | --- | --- |
| ***Signal transduction*** |  |  |  |
| *ARHGAP9 – COL4A4* | -0.51 | 0.28 | 0.79 |
| *ARHGAP9 – THBS3* | 0.39 | -0.48 | -0.87 |
| *CCL2 – COL4A4* | -0.95 | -0.07 | 0.88 |
| *CCL2 – CREB1* | 0.93 | -0.09 | -1.02 |
| *CCL2 – CRHR1* | 0.68 | -0.11 | -0.79 |
| *CCL2 – THBS3* | 0.68 | -0.19 | -0.87 |
| *CCNC – COL4A4* | 0.46 | -0.65 | -1.11 |
| *CCNC – CRHR1* | -0.58 | 0.65 | 1.23 |
| *CCNC – THBS3* | -0.41 | 0.34 | 0.75 |
| *CHN1 – CRHR1* | 0.43 | -0.59 | -1.02 |
| *CHN1 – HIF1A* | -0.36 | 0.34 | 0.70 |
| *COL4A4 – CREB1* | -0.93 | -0.03 | 0.90 |
| *COL4A4 – NFATC1* | -0.77 | 0.50 | 1.27 |
| *COL4A4 – PDPK1* | 0.41 | -0.50 | -0.91 |
| *CREB1 – CRHR1* | 0.80 | -0.03 | -0.83 |
| *CREB1 – THBS3* | 0.79 | 0.05 | -0.74 |
| *CRHR1 – KIDINS220* | 0.59 | -0.14 | -0.73 |
| *CRHR1 – NFATC1* | 0.83 | -0.47 | -1.30 |
| *CRHR1 – PDPK1* | -0.57 | 0.34 | 0.91 |
| *CRHR1 – YES1* | 0.40 | -0.71 | -1.11 |
| *CTNNBIP1 – NFATC1* | 0.07 | 0.85 | 0.78 |
| *KIDINS220 – PDPK1* | -0.74 | 0.30 | 1.04 |
| *KIDINS220 – SOS2* | -0.86 | 0.29 | 1.15 |
| *KIDINS220 – YES1* | 0.70 | 0.00 | -0.70 |
| *NFATC1 – THBS3* | 0.72 | -0.34 | -1.06 |
| *PDPK1 – SOS2* | 0.63 | -0.22 | -0.85 |
| *SOS2 – YES1* | -0.61 | 0.17 | 0.78 |
| ***Immune system*** |  |  |  |
| *ATF2 – NFATC1* | -0.03 | -0.75 | -0.72 |
| *BIRC2 – PDPK1* | -0.73 | 0.13 | 0.86 |
| *CREB1 – EIF4E2* | -0.94 | 0.07 | 1.01 |
| *CREB1 – IL7* | 0.91 | -0.03 | -0.94 |
| *CREB1 – UBR4* | 0.90 | -0.28 | -1.18 |
| *CREB1 – XAF1* | 0.71 | -0.03 | -0.74 |
| *EIF4E2 – IL7* | -0.87 | 0.10 | 0.97 |
| *EIF4E2 – NFATC1* | -0.87 | 0.46 | 1.33 |
| *EIF4E2 – UBR4* | -0.95 | 0.40 | 1.35 |
| *EIF4E2 – XAF1* | -0.72 | 0.10 | 0.82 |
| *IL7 – UBR4* | 0.88 | -0.34 | -1.22 |
| *NFATC1 – UBR4* | 0.85 | -0.18 | -1.03 |
| *PDPK1 – UBR4* | -0.55 | 0.34 | 0.89 |
| *UBA5 – UBR4* | 0.59 | -0.27 | -0.86 |
| *UBR4 – XAF1* | 0.88 | -0.16 | -1.04 |
| *UBR4 – YES1* | 0.35 | -0.59 | -0.94 |
| **Gene expression** |  |  |  |
| *AARS2 – ZNF253* | -0.57 | 0.23 | 0.80 |
| *AARS2 – ZNF606* | -0.91 | 0.21 | 1.12 |
| *AARS2 – ZNF711* | -0.60 | 0.31 | 0.91 |
| *RRN3 – ZNF184* | -0.51 | 0.34 | 0.85 |
| *RRN3 – ZNF606* | -0.06 | 0.87 | 0.93 |
| *RRN3 – ZNF711* | -0.22 | 0.50 | 0.72 |
| **Metabolism** |  |  |  |
| *ACSL4 – ARSK* | 0.56 | -0.66 | -1.22 |
| *ACSL4 – CCNC* | 0.75 | -0.74 | -1.49 |
| *ACSL4 – GPT* | -0.66 | 0.85 | 1.61 |
| *ACSL4 – NDUFA1* | 0.81 | 0.05 | -0.76 |
| *ACSL4 – PIKFYVE* | -0.22 | 0.69 | 0.91 |
| *ARSK – GM2A* | 0.48 | -0.34 | -0.82 |
| *CCNC – GM2A* | 0.65 | -0.44 | -1.09 |
| *CCNC – PSAT1* | 0.88 | -0.23 | -1.11 |
| *GPT – NDUFA1* | -0.59 | 0.24 | 0.83 |
| *GPT – PIKFYVE* | -0.05 | 0.79 | 0.84 |
| *GPT – PSAT1* | -0.65 | 0.46 | 1.11 |
| **Metabolism of proteins** |  |  |  |
| *ARSK – DPP4* | -0.27 | 0.57 | 0.84 |
| *CCL2 – DPP4* | 0.79 | -0.26 | -1.05 |
| *CCL2 – PCSK1* | 0.90 | -0.16 | -1.06 |
| *CCL2 – SPON2* | 0.77 | -0.03 | -0.80 |
| *DPP4 – GNE* | 0.74 | -0.21 | -0.95 |
| *DPP4 – MAGT1* | -0.57 | 0.37 | 0.94 |
| *MAGT1 – PCSK1* | -0.85 | 0.21 | 1.06 |
| *PCSK1 – SPON2* | 0.84 | 0.01 | -0.83 |
| **Developmental biology** |  |  |  |
| *CCNC – COL4A4* | 0.46 | -0.65 | -1.11 |
| *CLTB – SCN2B* | -0.39 | 0.37 | 0.76 |
| *COL4A4 – CREB1* | -0.93 | -0.03 | 0.90 |
| *SOS2 – YES1* | -0.61 | 0.17 | 0.78 |
| **Metabolism of lipids and lipoproteins** |  |  |  |
| *ACSL4 – ARSK* | 0.56 | -0.66 | -1.22 |
| *ACSL4 – CCNC* | 0.75 | -0.74 | -1.49 |
| *ACSL4 – PIKFYVE* | -0.22 | 0.69 | 0.91 |
| *ARSK – GM2A* | 0.48 | -0.34 | -0.82 |
| *CCNC – GM2A* | 0.65 | -0.44 | -1.09 |
| ***Axon guidance*** |  |  |  |
| *CLTB – SCN2B* | -0.39 | 0.37 | 0.76 |
| *COL4A4 – CREB1* | -0.93 | -0.03 | 0.90 |
| *SOS2 – YES1* | -0.61 | 0.17 | 0.78 |
| ***Innate immune system*** |  |  |  |
| *ATF2 – NFATC1* | -0.03 | -0.75 | -0.72 |
| *BIRC2 – PDPK1* | -0.73 | 0.13 | 0.86 |
| ***Disease*** |  |  |  |
| *CCNC – CHMP4C* | 0.60 | -0.38 | -0.98 |
| *CHMP4C – CREB1* | -0.85 | 0.31 | 1.16 |
| *CHMP4C – PDPK1* | 0.44 | -0.63 | -1.07 |

In a previous study by Taylor^49^, correlation was classified into three categories based on correlation coefficients: strong (0.68 ≤ CC ≤ 1.00), moderate (0.36 ≤ CC ≤ 0.67) and weak (CC ≤ 0.35). To increase the sensitivity of the assay for space travel, it is rational to only include gene pairs with correlation changing from strong to weak, weak to strong, or strong to opposite strong besides |ΔCC| > 0.70. Using this guideline, we identified the following gene pairs; 12 gene pairs (*CCL2* – *COL4A4, CCL2* – *CREB1, CCL2* – *CRHR1, CCL2* – *THBS3, COL4A4* – *CREB1, CREB1* – *CRHR1, CREB1* – *THBS3, CTNNBIP1* – *NFATC1, KIDINS220* – *PDPK1, KIDINS220* – *SOS2, KIDINS220* – *YES1*, and *NFATC1* – *THBS3*) from signal transduction, 11 gene pairs (*ATF2* – *NFATC1, BIRC2* – *PDPK1, CREB1* – *EIF4E2, CREB1* – IL7, *CREB1* – *UBR4, CREB1* – *XAF1, EIF4E2* – *IL7, EIF4E2* – *XAF1, IL7* – *UBR4, NFATC1* – *UBR4*, and *UBR4* – *XAF1*) from immune system, 2 gene pairs (*AARS2* – *ZNF606* and *RRN3* – *ZNF606*) from gene expression, 5 gene pairs (*ACSL4* – *CCNC, ACSL4* – *NDUFA1, ACSL4* – *PIKFYVE, CCNC* – *PSAT1* and *GPT* – *PIKFYVE*) from metabolism, 6 gene pairs (*CCL2* – *DPP4, CCL2* – *PCSK1, CCL2* – *SPON2, DPP4* – *GNE, MAGT1* – *PCSK1* and *PCSK1* – *SPON2*) from metabolism of proteins, 1 gene pair (*COL4A4* – *CREB1*) from developmental biology, 2 gene pairs (*ACSL4* – *CCNC* and *ACSL4* – *PIKFYVE*) from metabolism of lipids and lipoprotein, 1 gene pair (*COL4A4* – *CREB1*) from axon guidance, 2 gene pairs (*ATF2* – *NFATC1* and *BIRC2* – *PDPK1*) from innate immune system, and 1 gene pair (*CHMP4C* – *CREB1*) from disease. In total, 32 genes were selected, and they are *AARS2, ACSL4, ATF2, BIRC2, CCL2, CCNC, CHMP4C, COL4A4, CREB1, CRHR1, CTNNBIP1, DPP4, EIF4E2, GNE, GPT, IL7, KIDINS220, MAGT1, NDUFA1, NFATC1, PCSK1, PDPK1, PIKFYVE, PSAT1, RRN3, SOS2, SPON2, THBS3, UBR4, XAF1, YES1*, and *ZNF606*. To get a global view of these genes in regulating biological functions, we constructed the protein-protein interaction (PPI), disease-gene, drug-gene and miRNA-gene networks for the 218 DEGs and labelled the 32 genes in the networks (Figure 3). Genes *CCNC, CTNNBIP1, EIF4E2* and *RRN3* are not present in the disease-gene network; genes *BIRC2, CCNC, CHMP4C, CTNNBIP1, EIF4E2, IL7, KIDINS220, MAGT1, PIKFYVE, RRN3, SOS2, SPON2, THBS3, XAF1* and *ZNF606* are not present in the drug-gene network; and genes *CRHR1, DPP4, GPT* and *SPON2* are not present in the miRNA-gene network, respectively. Most of the 32 identified genes have high degrees of connectivity and are likely to be biological hubs. Thus, we propose that an RT^2^-PCR assay of these 32 genes could be a novel, effective and comparatively cheap way to continuously monitor astronauts’ health in space. Validation of this assay under microgravity simulation conditions is warranted for further optimization of the gene list before sending an assay kit for practical tests in space. Finally, we extracted the drug molecules (both approved and experimental, listed in Supplementary File 3: Drug_list.xls) and miRNAs (listed in Supplementary File 4: miRNA_list.xls), which potentially target these 32 genes from the drug-gene and miRNA-gene networks. These drugs and miRNAs could be applied as possible medical interventions for health conditions associated with space travel.

**Figure 3.**
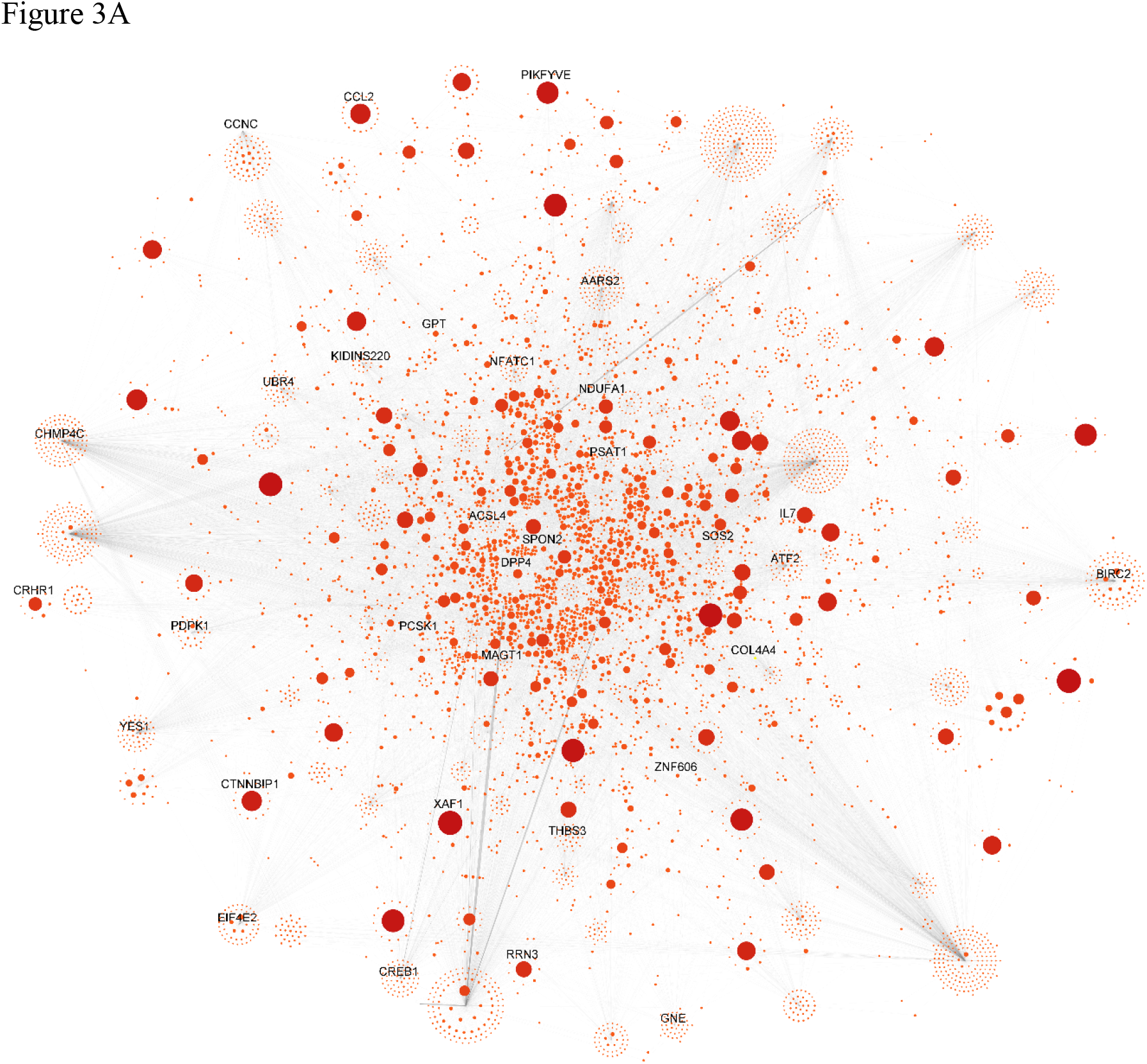
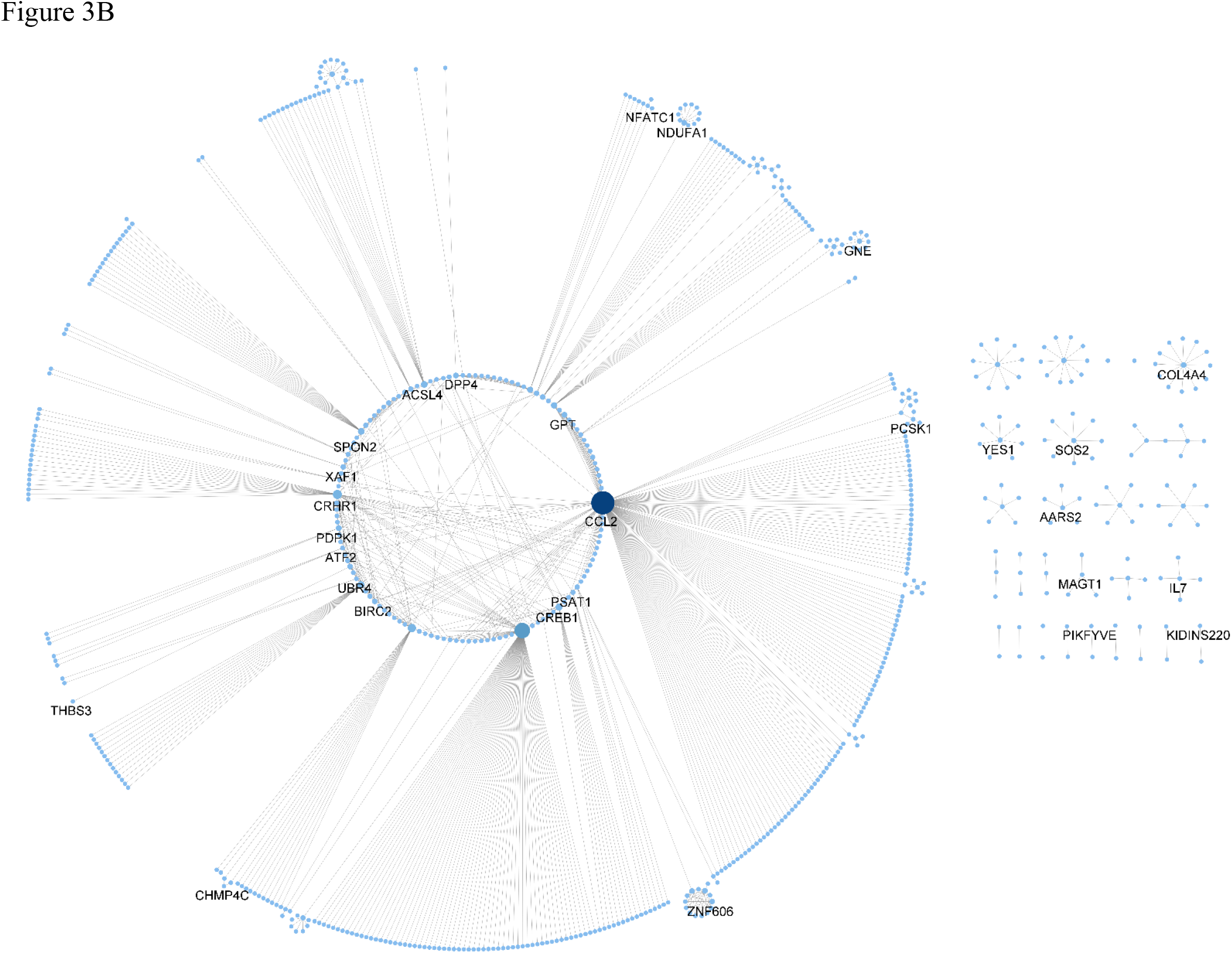
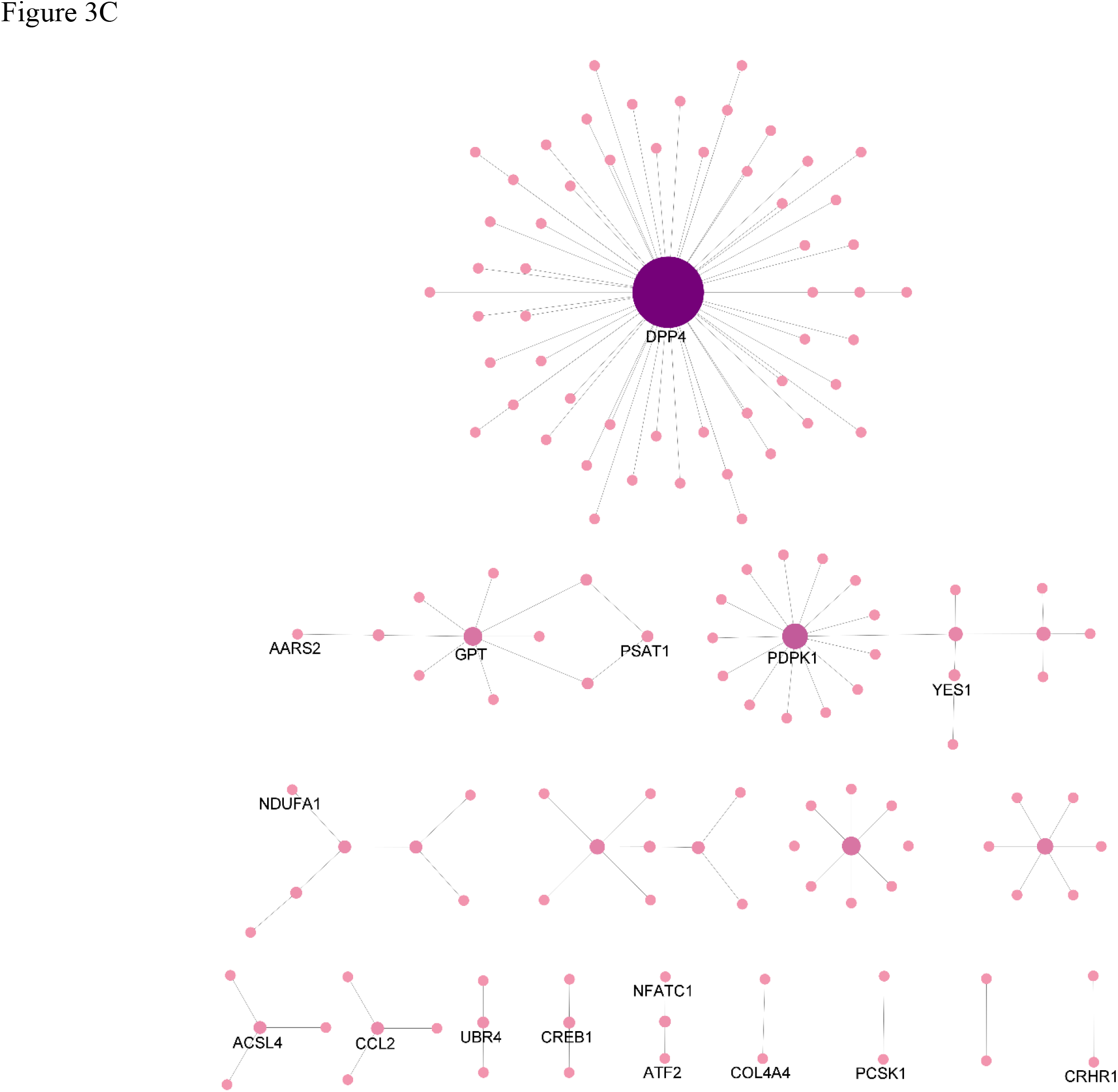
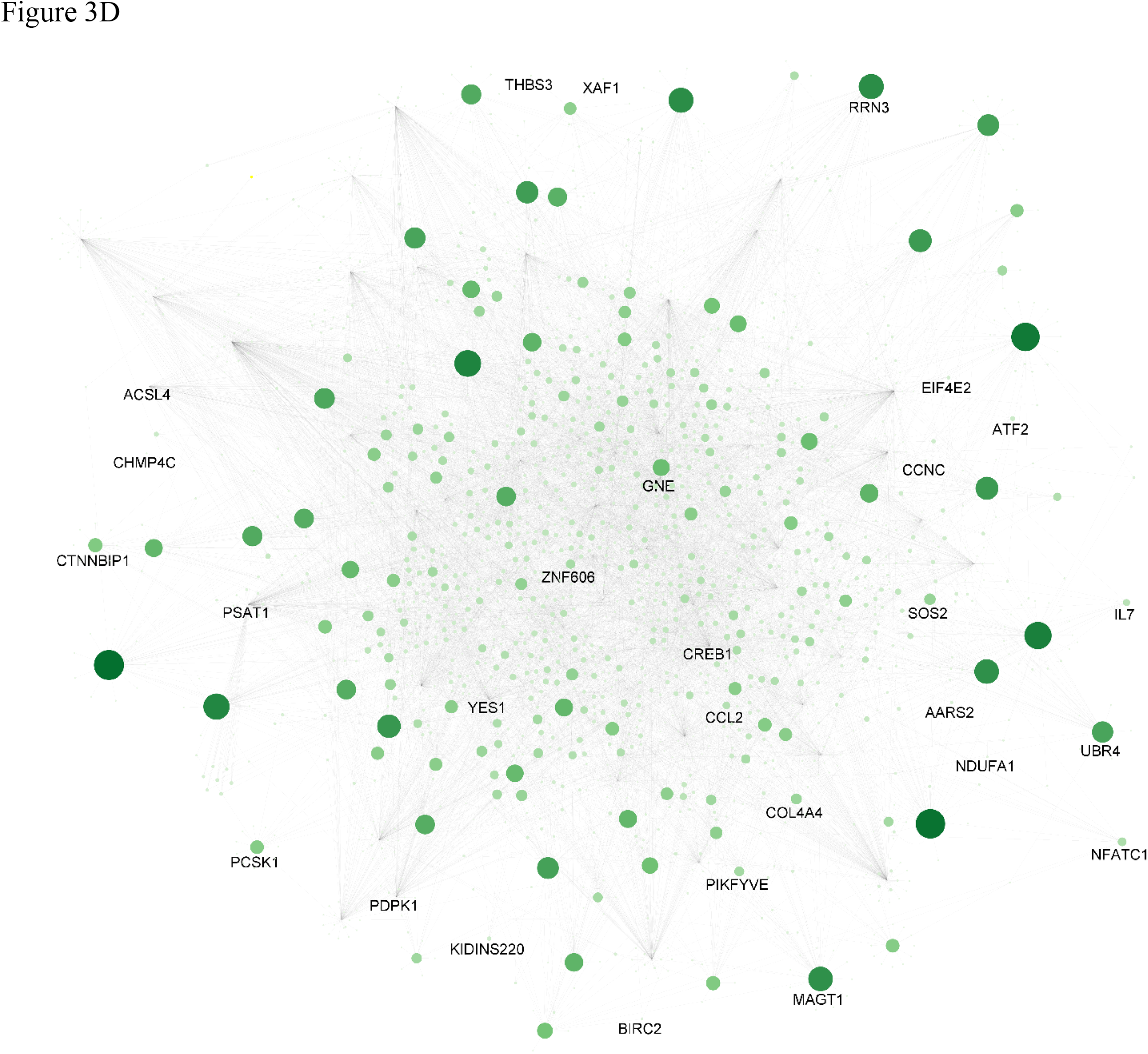
Protein-protein interaction (**A**), disease-gene (**B**), drug-gene (**C**) and miRNA-gene (**D**) networks for the 218 differentially expressed genes (DEGs) between Pre-flight and In-flight gene expression datasets collected from 10 astronauts (8 males and 2 females). Nodes with higher degrees of connectivity were shown in larger size and darker color. Genes identified for the rapid RT^2^-PCR assay kit were labelled in the networks. Genes *CCNC, CTNNBIP1, EIF4E2* and *RRN3* are not present in the disease-gene network; genes *BIRC2, CCNC, CHMP4C, CTNNBIP1, EIF4E2, IL7, KIDINS220, MAGT1, PIKFYVE, RRN3, SOS2, SPON2, THBS3, XAF1* and *ZNF606* are not present in the drug-gene network; and genes *CRHR1, DPP4, GPT* and *SPON2* are not present in the miRNA-gene network, respectively.

## Methods

### Data acquisition

Microarray dataset E-GEOD-74708, which is the largest publicly available space gene expression dataset, was downloaded from the NASA GeneLab Database (https:/genelab-data.ndc.nasa.gov/genelab). It contains gene expression profiles for 10 astronauts (8 male and 2 female), who stayed in the ISS between July 2009 and February 2013. For each astronaut, 2 profiles were collected at each of the three time points, Pre-flight (prior to departing for the ISS), In-flight (while staying at the ISS) and Post-flight (after return to the ground).

### Data processing

Individual gene expression profiles were combined into three DataFrames: Pre-flight, In-flight, and Post-flight using the Pandas package (<u>version</u> 1.3.0)^50^ in Python. These DataFrames were analyzed for possible outliers and verified for accuracy. Gene expression readings with null values were removed. Then, Agilent IDs in the data were mapped to gene names using the Ensembl BioMart database (https://www.ensembl.org/biomart/martview).

### DEG identification and signaling pathway analysis

Differentially expressed genes (DEGs: |log_2_FC| ≥ 3.00 and *p* < 0.05) were identified between the In-flight and the Pre-flight DataFrames using the DEGseq package (version 1.42.0) in R^51^. The Likelihood Ratio Test (LRT) function was applied in the calculation. No DEGs were identified between the Post-flight and the In-flight DataFrames. The DEGs between In-flight and Pre-flight were then subjected to signaling pathway analysis using the InnateDB tool^52^.

### Calculation of gene pair correlation matrices

Pairwise Pearson correlation coefficient matrices for the 3 gene expression DataFrames were calculated using formula 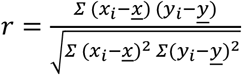. The correlation matrices were subsequently transferred to be visualized using an R platform. To analyze gene pair correlations for the signaling pathways, the expression profiles of the genes for the pathways were extracted from the DataFrames to produce the DataFrame subsets. For each signalling pathway, pairwise Pearson correlation coefficients ware calculated and visualized as outlined above between the In-flight and Pre-flight subsets and between the Post-flight and In-flight subsets. Positive and negative correlations were represented in blue and red, respectively. Additionally, gene pair correlation coefficient differentials, ΔCC (In-Pre) = CC_In-flight_ – CC_Pre-flight_ and ΔCC (Post-In) = CC_Post-flight_ – CC_In-flight_ were calculated by matrix subtraction between the In-flight and Pre-flight subsets and between the Post-flight and In-flight subsets. These were subsequently visualized with correlation difference plots with positive and negative ΔCCs shown in green and red, respectively.

### Protein-protein interaction network and disease network constructions

The BioGrid database^53^ and DisGeNET database^54^ were downloaded in March 2021 to create a protein-protein interaction (PPI) network (Figure 3A) and extract the disease interaction information (Figure 3B), respectively, for the DEGs. The networks were then constructed using Python and visualized using Cytoscape^55^.

### Drug and miRNA screenings

The Drugbank^56^ and miRTARBase^57^ databases were also downloaded in March 2021 to extract the drug (Figure 3C) and miRNA (Figure 3D) interaction information for the DEGs. The networks were subsequently constructed using Python and visualized using Cytoscape^55^. From these two networks, we identified potential drugs and miRNAs targeting the 32 genes we identified to be used in a rapid RT^2^-PCR assay for space travel (Supplementary File 3: Drug_list.xls and Supplementary File 4: miRNA_list.xls).

## Conclusion

In this study, we analyzed the effects of space travel on gene expression and correlation using the publicly available microarray dataset E-GEOD-74708 and systematically developed a novel rapid assay which could be used to monitor astronauts’ health in space. However, this study faces several limitations. First, the sample size (10 astronauts) is small due to the nature of space travel and unstandardized astrobiological research. More sample collection is needed to optimize the monitoring assay. Secondly, there were only 2 female astronauts in the microarray dataset. To get better representation and more reliable accuracy in the analysis, more gene expression profiles from a diverse population of astronauts should be obtained. Thirdly, our laboratory is not equipped with any microgravity infrastructure, and thus, we cannot validate the proposed assay. However, in our future studies, we will reach out to resources such as microgravity simulators and continue to conduct research in optimizing and validating this assay. Despite these drawbacks, our current study proposes a new strategy to develop genome-based rapid assays. This approach could also be applied to other research fields, such as cancer diagnostic assays.

## Author contribution

This research was originated by J.Y. and carried out by A.S. and J.Y.

